# Methionine synthase supports tumor tetrahydrofolate pools

**DOI:** 10.1101/2020.09.05.284521

**Authors:** Joshua Z. Wang, Jonathan M. Ghergurovich, Lifeng Yang, Joshua D. Rabinowitz

**Author notes:** These authors contributed equally to this work (Joshua Z. Wang, Jonathan M. Ghergurovich). Corresponding author: Joshua Rabinowitz, Department of Chemistry and the Lewis-Sigler Institute for Integrative Genomics, Princeton University, Washington Rd, Princeton, NJ 08544, USA.

## Abstract

Mammalian cells require activated folates to generate nucleotides for growth and division. The most abundant circulating folate species is 5-methyl tetrahydrofolate (5-methyl-THF), which is used to synthesize methionine from homocysteine via the cobalamin-dependent enzyme methionine synthase (MTR). Cobalamin deficiency traps folates as 5-methyl-THF. Here, we show using isotope tracing that methionine synthase is only a minor source of methionine in cell culture, tissues, or xenografted tumors. Instead, methionine synthase is required for cells to avoid folate trapping and assimilate 5-methyl-THF into other folate species. Under conditions of physiological extracellular folates, genetic MTR knockout in tumor cells leads to folate trapping, purine synthesis stalling, nucleotide depletion, and impaired growth in cell culture and as xenografts. These defects are rescued by free folate but not one-carbon unit supplementation. Thus, MTR plays a crucial role in liberating tetrahydrofolate for use in one-carbon metabolism.

## Introduction

Dividing cells require activated one-carbon (1C) units to synthesize and package nucleic acids [1]. In mammals, tetrahydrofolate (THF) species carry the formyl- and methylene-1C groups necessary for purine and thymidylate biosynthesis [2–4]. This essentiality is reflected in the efficacy of antifolate chemotherapeutic agents for treating malignancy, their associated toxicities in proliferating normal tissues, and alleviation of these effects by folate rescue agents (i.e. 5-formyl-THF, a.k.a. leucovorin) [5]. The 1C units of activated folates are locally sourced from serine, glycine, and other amino acids [6, 7], with the major 1C donor typically serine via the enzyme serine hydroxymethyltransferase (SHMT).

In contrast to the capacity to manipulate 1C units using THF, mammals lack the necessary machinery for *de novo* folate synthesis. Folate species are instead absorbed from food and converted in the gut and liver to 5-methyl tetrahydrofolate (5-methyl-THF), the dominant circulating folate species, before distributing to tissue folate pools [8, 9]. Deficiency in dietary folate causes neural tube defects during pregnancy [10].

Mammals also lack the ability to assimilate inorganic sulfur, and thus have a daily requirement for sulfur containing amino acids like methionine [11–13]. In addition to its role in protein synthesis, methionine serves as the primary methyl source for DNA, protein, and lipid methylation via S-adenosyl-methionine (SAM) [14]. Methyl donation by SAM ultimately yields the free thiol species homocysteine, which undergoes either degradation via the transsulfuration pathway to yield cysteine or remethylation to methionine [11]. SAM also acts as an allosteric regulator, inhibiting methylenetetrahydrofolate reductase (MTHFR), the enzyme producing 5-methyl-THF, and activating cystathionine β-synthase (CBS), the first enzyme of transsulfuration [15, 16]. Low SAM accordingly tends to preserve homocysteine and promote its conversion to methionine, allowing homocysteine to substitute for dietary methionine [17–19].

Methionine synthase (MTR) catalyzes the methyl transfer from 5-methyl-THF to homocysteine, yielding methionine and tetrahydrofolate (THF), linking folate and methionine cycling [11, 20] (Figure 1*A*). MTR is one of two enzymes in humans known to utilize the cofactor vitamin B_12_ (cobalamin) [21]. Mutations in *MTR* leading to diminished enzyme activity result in methylcobalamin deficiency G (cblG) disorder, a condition characterized by hyperhomocysteinemia, hypomethioninemia and homocystinuria [22]. In development, *Mtr* is essential, leading to embryonic death in mice [23]. MTR is universally expressed in tissues, while expression of betaine-homocysteine S-methyltransferase (BHMT), the other enzyme known to produce methionine via methyl transfer from betaine to homocysteine, is primarily expressed in the liver and, to a lesser degree, in the kidney ([15, 24–26]; Figure 1*B*). Similarly, MTR is broadly expressed in human cancer cell lines [27] and tumors [28].

**Figure 1.**
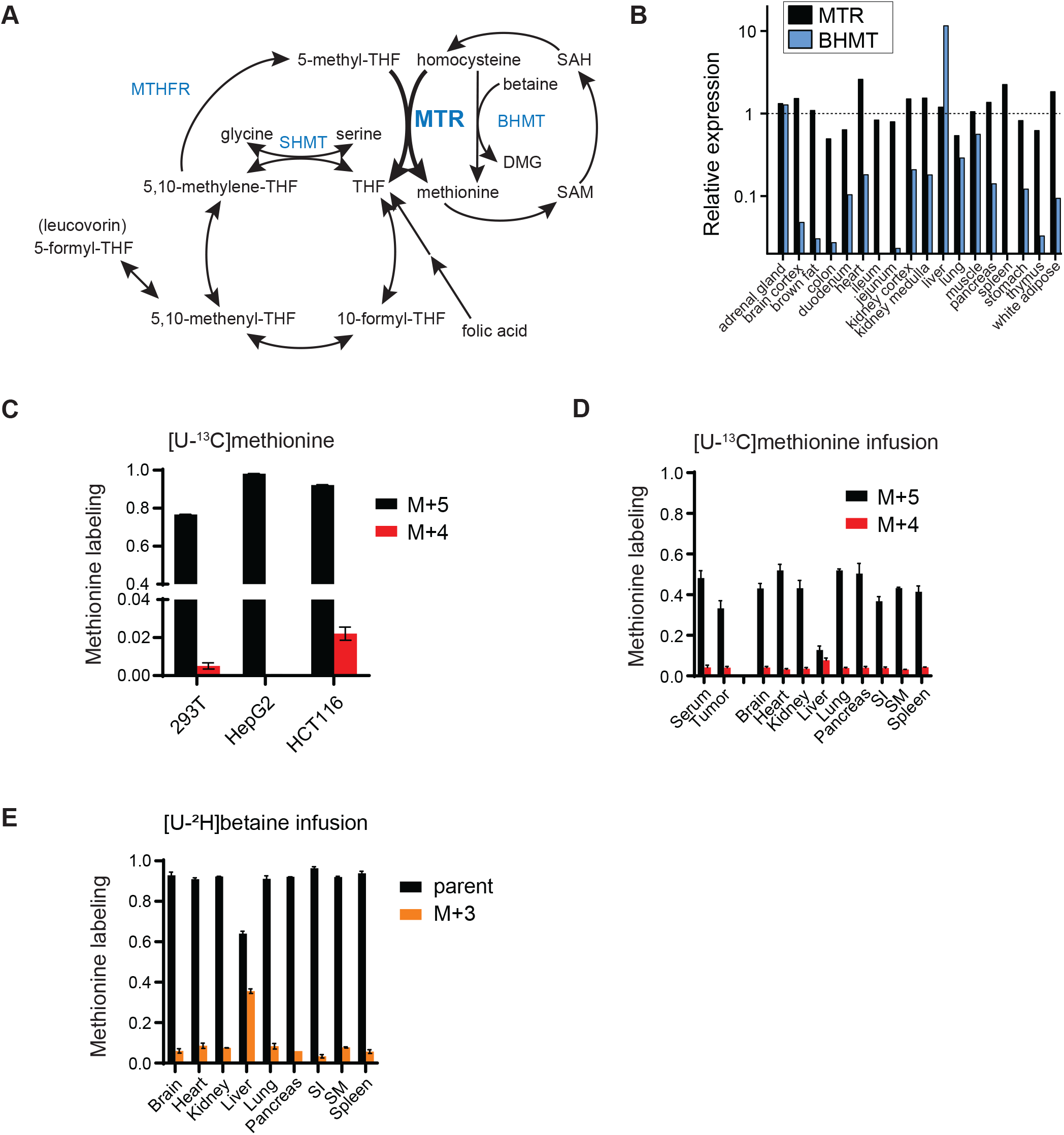
MTR is a minor source of methionine in cell lines, tissues and tumors. *(A)* Schematic showing the activity of MTR at the intersection of the folate and methionine cycles. MTR = methionine synthase, MTHFR = methylenetetrahydrofolate reductase, BHMT = betaine-homocysteine S-methyltransferase, SHMT = serine hydroxymethyltransferase. *(B)* Relative protein levels of MTR and BHMT in mice from [24]. *(C)* Methionine labeling in cell lines after culturing for 4 h in media containing [U-^13^C]methionine (mean ± SD, n = 2 for HCT116 cells, 3 for 293T and HepG2). *(D)* Methionine labeling of serum, PDAC tumors, and normal tissues in C57BL/6J mice after [U-^13^C]methionine infusion for 2.5 h (mean ± SD, n=3 mice; two technical replicates were included for each tumor). (E) Methionine labeling in normal tissues in C57BL/6J mice after [U-^2^H]betaine infusion for 4 h (mean ± SD, n=1 infusion; two technical replicates for each tissue).

The physiological role of MTR is evident from vitamin B_12_ deficiency. Classically, B12-deficient patients present with neurological and hematological manifestations, most notably a macrocytic, megaloblastic anemia, an indicator of impaired nucleotide synthesis. This phenotype similarly occurs in folate deficiency [29, 30]. vitamin B_12_ restriction or chemical inactivation of vitamin B_12_ (and thus MTR) by nitrous oxide impairs MTR activity *in vitro* and *in vivo* [30–32]. Moreover, because methylene-THF reduction to 5-methyl-THF is nearly irreversible in cells (due to high cytosolic NADPH/NADP^+^) [33], MTR dysfunction leads to folate trapping as 5-methyl-THF [20, 30, 34–36].

Despite numerous studies over the last 70 years into folate and methionine metabolism, it is unclear how much methionine arises locally from remethylation in a given tissue. Similarly, the role of MTR is supporting cancer growth has not yet been robustly examined. Here, we combine modern genetic tools with isotope tracer studies to dissect the role of MTR in normal tissue and tumor biology. We show that tissue and tumor methionine is overwhelmingly acquired from the systemic circulation, with MTR contributing only a minor fraction to local methionine pools. Conversely, MTR plays a major role in maintaining folate pools, with cells and xenografted tumors lacking MTR unable to grow when subjected to physiological folate conditions. Mechanistically, we show that MTR loss leads to folate trapping, purine synthesis stalling and nucleotide depletion, which can be mitigated with folate rescue agents.

## Results

### Methionine comes mainly from uptake rather than homocysteine remethylation

Methionine cycling involves reduction of 5,10-methylene-THF to 5-methyl-THF mediated by MTHFR, followed by conversion of homocysteine to methionine by MTR. Alternatively, homocysteine can be converted to methionine by betaine hydroxymethyltransferase (BHMT) (Figure 1*A*). MTR is expressed throughout the body, while BHMT is preferentially expressed in liver (Figure 1*B*).

To probe methionine cycling, we began with cell culture experiments where the media methionine component was substituted for [U-^13^C]methionine at the same concentration. We reasoned that untransformed methionine would retain all five labeled carbons (M+5). In contrast, methionine that had been recycled via homocysteine remethylation would lose one ^13^C carbon, retaining four labeled carbons (M+4; Figure 1*A*, Supplementary figure S1*A*). The fraction of M+4 to M+5 methionine approximates the relative amount of cellular methionine arising via remethylation as opposed to uptake. All cell lines examined displayed a high fraction of M+5 and low fraction of M+4 methionine (<3%), consistent with intracellular methionine mainly arising from uptake (Figure 1*C*). Similar experiments with either [U-^13^C]- or [3-^13^C]serine tracing, looking for M+1 methionine coming from homocysteine remethylation via the SHMT-MTHFR-MTR pathway (Supplementary figure S1*B*) showed very low levels of enrichment (<1% enrichment) (Supplementary figure S1*C*). Thus, consistent with prior literature, methionine cycling is minimal in cell culture[3, 37].

To examine whether these observations hold in vivo, we infused [U-^13^C]methionine via jugular venous catheter into C57BL/6J mice bearing subcutaneous pancreatic ductal adenocarcinoma (PDAC) allograft tumors [38], achieving an enrichment of 30-60% in serum, tumors and most tissues (Figure 1*D*). This extent of labeling is adequate for robust label detection while avoiding severe metabolic perturbation by the labeled methionine infusion. In all tissues except liver, M+5 labeling was dominant, with M+4/M+5 ratio about 0.1, suggesting that approximately 10% of methionine arose from homocysteine remethylation (Figure 1*D*). The fraction of M+4 was similar between serum and these tissues, and thus does not necessarily reflect tumor-intrinsic methionine cycling. In contrast, liver M+5 labeling was markedly lower and M+4 higher, resulting in a M+4/M+5 ratio of ~0.6 and consistent with substantial methionine arising via homocysteine remethylation. However, infusion of labeled serine demonstrated limited folate-mediated methionine synthesis throughout the body, including the liver (Supplementary figure S1*E*). Thus, homocysteine remethylation in the liver must utilize a different methyl donor.

We next sought to assess the contribution of remethylated methionine from betaine, the alternative source of methionine in mammals. The BHMT reaction transfers a methyl group from the quaternary amine of betaine to homocysteine via a cobalamin-independent reaction mechanism [39]. Mice were infused with the [U-^2^H]betaine tracer, which contains three deuterium (^2^H) on each methyl group and thus will give rise to M+3 labeled methionine (Supplementary figure 1*F*). This infusion revealed preferential methionine synthesis from betaine in liver (Figure 1*E*, Supplementary figure 1*G*). Collectively, these studies support BHMT being a source of liver methionine while the MTHFR-MTR pathway operates more broadly throughout the body but, at least acutely, makes only a small contribution to methionine overall.

### MTR is required for cell growth using physiological folates

The above studies raised questions about the physiological role of the MTHFR-MTR pathway, since it was not a dominant methionine producer anywhere in the body. As an alternative strategy to examine the role of MTR, we generated clonal MTR knockout cell lines (ΔMTR) using CRISPR-Cas9 gene editing in three cancer cell backgrounds: HCT-116 (colorectal carcinoma), 8988T (pancreatic ductal carcinoma), and HepG2 (hepatocellular carcinoma) (Figure 2*A*). Each of these parental cell lines possesses reasonable expression of MTR (Supplementary figure S2*A*). In standard tissue culture media formulations containing folic acid, each of these MTR knockout lines grew reasonably well, albeit at slower rates than control cells in the 8988T and HepG2 backgrounds (Figure 2*B*). Expectedly, neither MTR knockout or control lines grew in the complete absence of folates (Supplementary figure S2*B*). Control cells were, however, able to grow rapidly in media where folic acid was replaced with 5-methyl-THF, the most abundant circulating folate species. In contrast, MTR knockout cells did not grow at all (Figure 2*B*). The defective growth of the MTR knockout cells was fully rescued through supplementation with 5-formyl-THF (leucovorin) (Figure 2*B*), a folate species that enters folate metabolism independently of MTR. Thus, MTR is required for cancer cell growth using 5-methyl-THF, the main circulating folate species, presumably because MTR is required to liberate THF from 5-methyl-THF.

**Figure 2.**
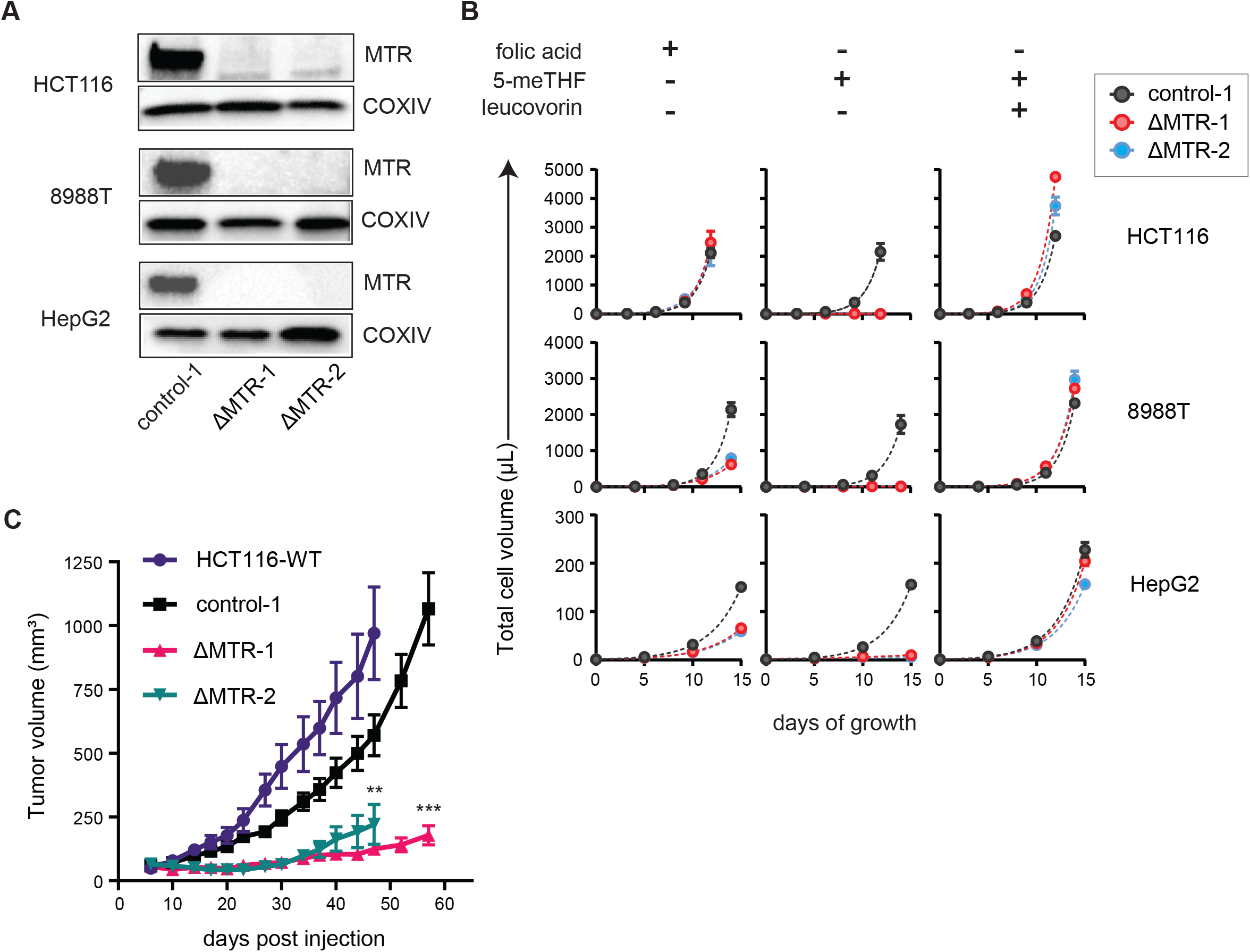
MTR is essential for cell and tumor growth under physiological conditions. *(A)* Western blot analysis of MTR in HCT116, 8988T and HepG2 clonally isolated lines after treatment with lentiviral CRISPR-Cas9 targeting scrambled control (control-1) or MTR (ΔMTR-1 and ΔMTR-2). *(B)* Cell growth curves in the media containing indicated folate sources (mean ± SD, n=2). 5-meTHF = 5-methyl-tetrahydrofolate. *(C)* Growth of subcutaneous HCT116 xenografts (mean ± SEM, n=10 mice, Student’s t-test). ** = p < 0.01, *** = p < 0.001 for comparison to respective controls. WT = wild-type.

In addition to 5-methyl-THF, other species such as folic acid and 5-formyl-THF are present in serum at lower concentrations [40]. To investigate whether MTR is required for tumor growth *in vivo*, we implanted HCT116 ΔMTR cells subcutaneously into the lateral flanks of nude mice and monitored their growth. Tumors derived from MTR knockout cells demonstrated a growth deficiency (Figure 2*C*, Supplementary figure S2*C-D*), implicating MTR as being important for folate access in the presence of the physiological mixture of circulating folates.

### Loss of MTR depletes cells and tumors of reduced folate species

These above findings support the hypothesis that MTR is required to supply cancer cells with free THF when fed 5-methyl-THF, the most abundant physiologically relevant folate species. To evaluate this further, we measured cellular folate species in HCT116 cells cultured with either folic acid or 5-methyl-THF as the sole media folate source. Expectedly, MTR loss led to a significant buildup of 5-methyl-THF in both media conditions (Figure 3*A*), in line with the concept of “methyl trapping.” Concurrently, most downstream folate species were substantially decreased. The most depleted species was 5,10-methylene-THF, which is required for thymidine biosynthesis. In contrast, 10-formyl-THF, which is used for purine synthesis, was not strongly depleted. This likely reflects inhibition of 10-formyl-THF consumption by purine pathway disruption due to overall folate species dysregulation. Interestingly, buildup of 5-methyl-THF and depletion of other folate species was triggered by MTR loss in cells growing in media containing folic acid, indicating the importance of MTR for folate pool homeostasis even when adequate free folate is available.

**Figure 3.**
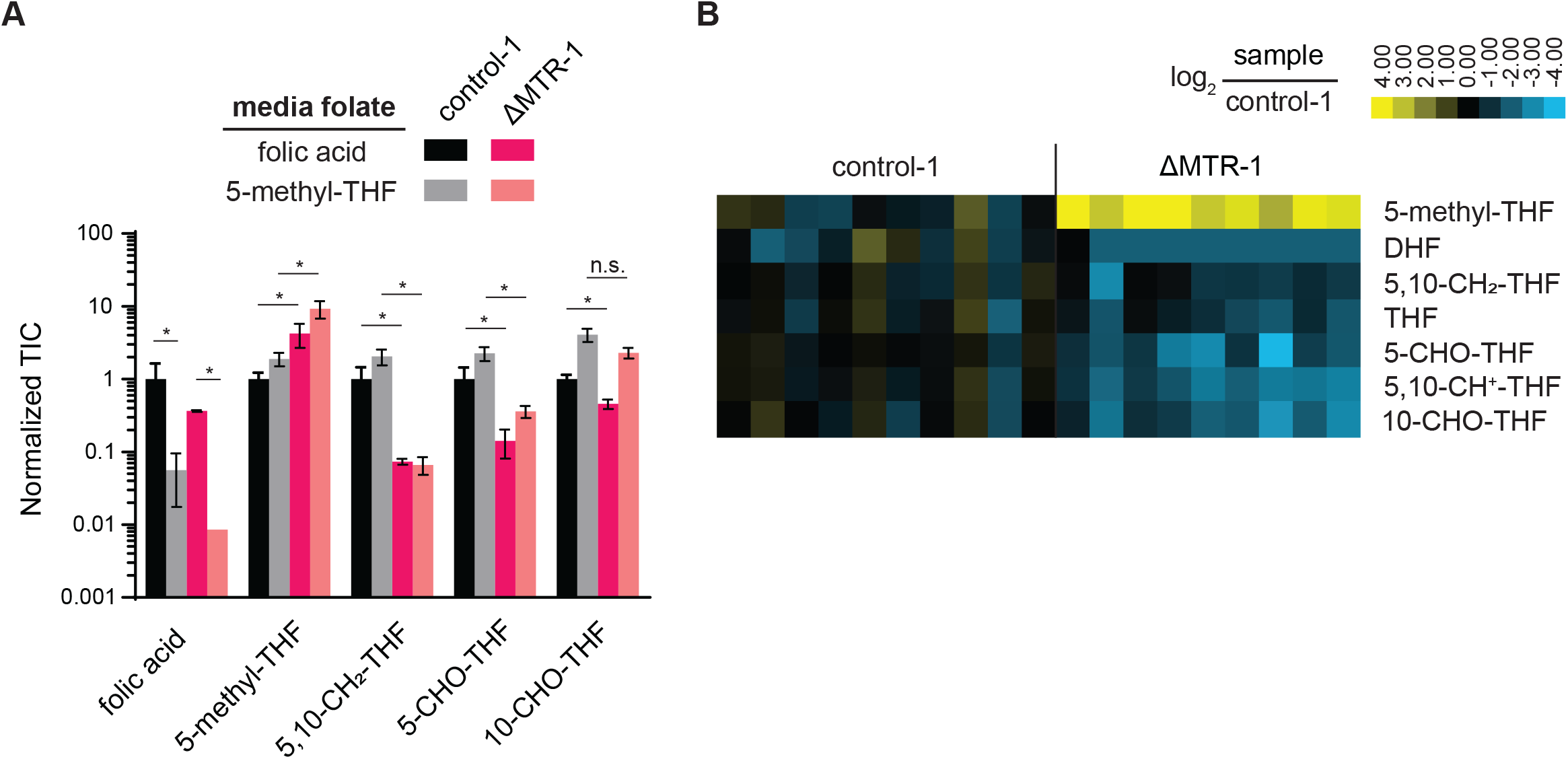
Loss of MTR results in the elevation of 5-methyl-THF and depletion of other folate species. *(A)* Relative abundance of folate species in HCT116 control and MTR knockout cells in culture across media with indicated folate sources. Intensities are normalized to the average of control-1 cells in folic acid (mean ± SD, n=2, one-way ANOVA, p-values compare control to MTR knockout in the same medium). *(B)* Relative abundance of folate species in HCT116 control and MTR knockout subcutaneous tumors. For each condition, individual biological replicates are shown, normalized to control tumors and analyzed in parallel by LC-MS. DHF = dihydrofolate, THF = tetrahydrofolate. TIC = total ion count, * = p < 0.05, n.s. = not significant.

In vivo, 5-methyl-THF was elevated ~12-fold in HCT116 ΔMTR-1 tumors when compared to control-1 tumors, while all other folate species were depleted by about 50-80% (Figure 3*B*). Thus, MTR is required to maintain folate pools under physiological folate conditions.

### MTR activity prevents stalling of purine and pyrimidine biosynthesis

We next sought to investigate the downstream effects of MTR loss and associated changes in folate species concentrations on other aspects of cellular metabolism. To this end, HCT116 ΔMTR cells and controls were cultured in media containing either folic acid or 5-methyl-THF as the sole media folate. Metabolomics demonstrated global changes in metabolism due to MTR loss (Supplementary figure S3*A*). Most notably, we observed dramatic increases in several purine biosynthetic intermediate pools, including GAR, SAICAR and AICAR (Figure 4*A*), and corresponding decreases in nucleotide species (Figure 4*B*, Supplementary figure S3*B*). Nucleotide depletion was most substantial in media containing 5-methyl-THF as the sole folate source. Similarly, MTR loss in folic acid media led to a buildup of dUMP, consistent with impaired thymidine biosynthesis secondary to diminished 5,10-methylene-THF, though the thymidylate species dTMP and dTTP were unperturbed (Supplementary figure S3*C*). Interestingly, both dUMP and dTMP/dTTP were significantly decreased in 5-methyl-THF media, possibly reflecting the overall decrease in the precursor UMP, perhaps due to purine-pyrimidine crosstalk.

**Figure 4.**
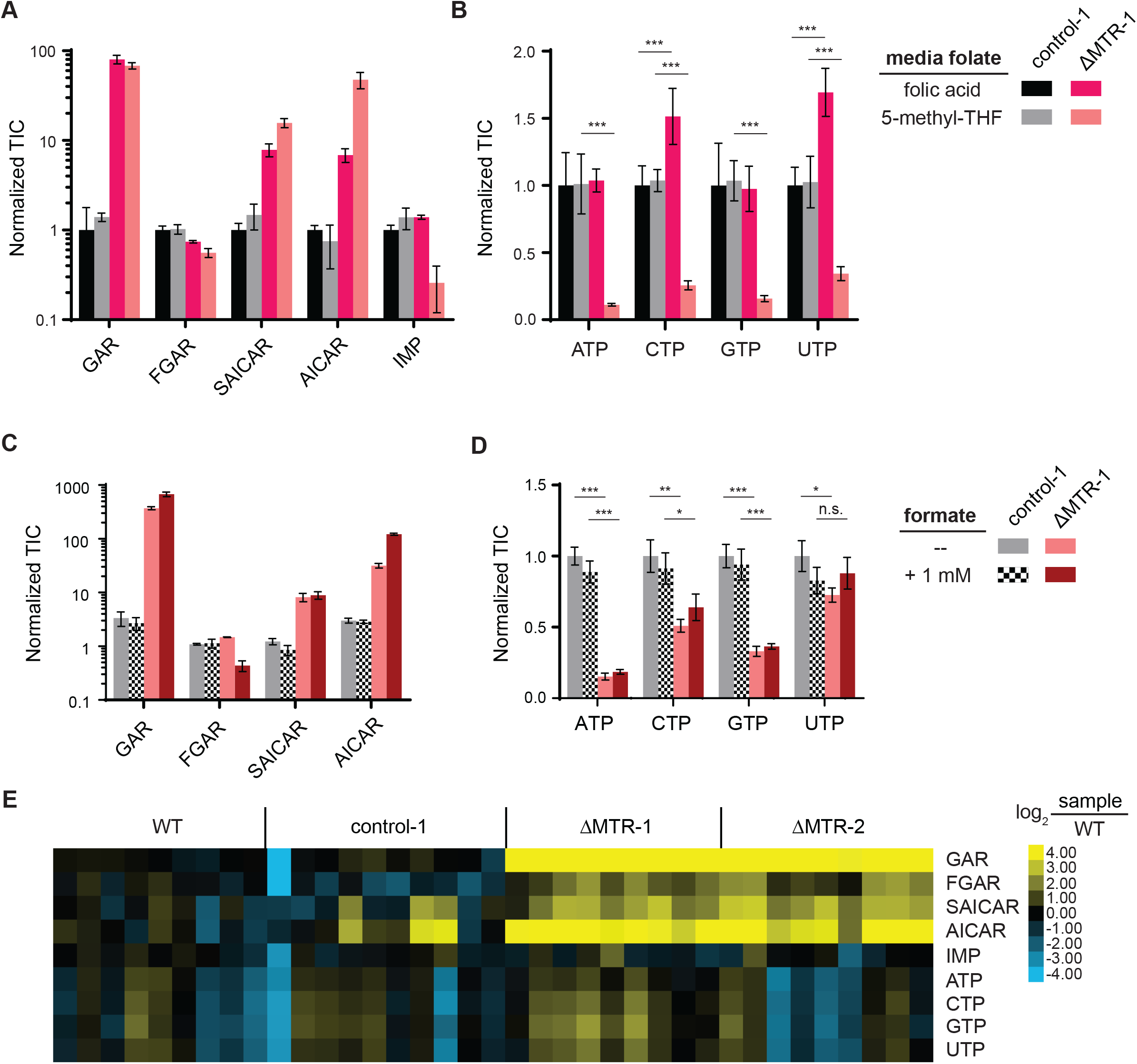
MTR supports purine biosynthesis by promoting folate availability. Relative abundance of *(A)* purine biosynthetic intermediates and *(B)* nucleotides in HCT116 control and MTR knockout cells in culture with indicated folate sources. Intensities are normalized to the average of control-1 cells in folic acid (mean ± SD, n=3). Relative abundance of *(C)* purine biosynthetic intermediate and *(D)* nucleotides in control and MTR knockout cells grown in media with 5-methyl-THF, with and without the addition of 1 mM formate (mean ± SD, n=3). *(E)* Relative abundance of purine biosynthetic intermediate and nucleotide triphosphates in HCT116 control and MTR knockout subcutaneous tumors. For each condition, individual biological replicates are shown, normalized to wild-type tumors and analyzed in parallel by LC-MS. TIC = total ion count, * = p < 0.05, ** = p < 0.01, *** = p < 0.001, n.s. = not significant. WT = wild-type.

To evaluate whether these changes were due to aberrant levels of folates, or deficiency of usable 1C units [3], we attempted to rescue these effects through supplementation with the 1C precursor formate. Consistent with this phenotype arising from folate dysregulation, formate did not rescue (Figure 4*C-D*).

To investigate whether these metabolic changes occur in vivo, we examined nucleotide levels from the HCT116 xenografts (Supplementary figure S4*A*). Elevated levels of GAR (50-fold), FGAR (4-fold), SAICAR (4-fold), and AICAR (7-fold) were observed in the MTR knockouts relative to wild-type controls, mirroring the phenotype observed in cell culture (Figure 4*E*). In contrast, however, IMP and nucleotides were not consistently depleted in tumors derived from ΔMTR knockouts (Figure 4*E*, Supplementary figure S4*A*), suggesting that tumors in vivo have other means of maintaining nucleotide homeostasis, perhaps through import of circulating nucleosides. Overall, these results demonstrate that MTR is required for folate species and nucleotide homeostasis under physiological folate conditions.

## Discussion

Folate metabolism has long been recognized as a central player in supporting proliferation in tissues and tumors. However, given its complexity, many questions remain pertaining to folate species homeostasis and how dysregulation of folate levels perturbs other areas of cellular metabolism. In this work, we show that despite its name, methionine synthase (MTR) is a minor immediate source of tissue and tumor methionine, but instead is acutely critical for maintaining folate pools when cells are exposed to physiological folate conditions. Furthermore, we find that MTR loss significantly impairs thymidine and purine biosynthesis, effects not rescued with 1C unit supplementation.

The essentiality of MTR depends on the available extracellular folate species, and its importance is lost when culturing cells using a standard (i.e. folic acid) media preparation. This reflects culture medium typically containing markedly supraphysiological levels of free folate, and lacking physiological levels of 5-methyl-THF. Notably, while there has been recent progress towards more physiological culture medium formulations [41, 42], these continue to employ vitamin levels, including folates, selected for convenience rather than physiological relevance. Future physiological medium should switch to 5-methyl-THF as the primary folate source, and more generally push towards physiological vitamin abundances.

The importance of MTR for growth using 5-methyl-THF as the folate source makes sense based on the pathway architecture. As expected for folate perturbations, intermediates accumulated in both the thymidine and purine pathways. But the ties between the observed changes in folate pools and nucleotide alterations was not obvious. Specifically, purine metabolism was dramatically perturbed by MTR loss, even though its direct folate substrate, 10-formyl-THF, was only modestly affected. Conversely, 5,10-methylene-THF was the most depleted folate species but thymidine levels were maintained. Why the stronger effects on purine metabolism? One possibility is that the structural similarity of different folate cofactors leads to enzyme inhibition by elevated species acting as competitive active site inhibitors. For example, elevated DHF arising from methotrexate treatment is known to inhibit multiple enzymes in folate and purine metabolism [43–45]. Similarly, 5-formyl-THF, and to a lesser extent 5-methyl-THF, have been shown to bind SHMT [46]. In the context of MTR loss, the most likely disrupter of purine synthesis is elevation of 5-methyl-THF. This possibility is supported by the observation that media supplemented with 5-methyl-THF exacerbates purine depletion (Figure 4*B*, Supplementary figure S4*B*). Interestingly, recent studies in MTR-null fibroblasts demonstrated buildup of 5-methyl-THF leads to impaired nuclear synthesis of thymidylate, but not purines [34]. However, these cells were cultured in media containing 5-formyl-THF, rather than 5-methyl-THF, bypassing the additional stress imposed by 5-methyl-THF media exposure. Though it remains unproven which enzymatic steps of purine synthesis are impacted by 5-methyl-THF, the excessive buildup of purine intermediates upstream of AICAR (Figure 4*A,C,E*) suggests inhibition of AICAR transformylase, the enzyme catalyzing the conversion of AICAR to IMP.

Is there a therapeutic role for MTR inhibitors in the treatment of cancer? Given the substantial success of antifolate treatments in combatting malignancy and MTR’s role in maintaining tumor folate pools, targeting MTR, either with a single agent or in combination with other antifolates, merits investigation[47]. Antifolate resistance presents a significant challenge for achieving remission [48, 49], and MTR inhibitors could potentially help in this regard by providing an orthogonal approach to depleting tumor folates and/or by promoting toxic 5-methyl-THF accumulation within tumor cells resistant to other antifolate agents. Side effects could potentially be mitigated by rescue therapy, perhaps with leucovorin, as is common for other anti-folates. To our knowledge, no clinically approved antifolate therapies are known to directly inhibit MTR, though some low potency, folate-like small molecule inhibitors have been reported [50–52]. We look forward to the advancement of such compounds into in vivo tools so that the therapeutic potential of MTR inhibition can be more rigorously evaluated.

## Materials and Methods

### Cell lines, culture conditions and reagents

HCT116, 293T, and HepG2 were obtained from ATCC (Manassas, VA). 8988T cells were obtained from DSMZ (Braunschweig, Germany). Unless otherwise stated, cells were maintained in DMEM (CellGro 10-017, Mediatech) supplemented with 10% fetal bovine serum (FBS, F2442, Sigma-Aldrich). Cell lines were cultured in a 37^°^C incubator with 5% CO_2_ and routinely screened for mycoplasma contamination by PCR. Rowell Park Memorial Institute medium (RPMI, Gibco 11875093; Thermo Fisher) supplemented with 20% FBS (v/v) was used for single-cell plating MTR knockout cells. Folic acid (BP25195) and leucovorin (18-603-788) were obtained from Fisher Scientific; 5-methyl-THF was obtained from Sigma Aldrich (M0132). Isotopic tracers [U-^13^C] methionine (CLM-893-H), [U-^13^C]serine (CLM-1574-H), [3-^13^C] serine (CLM-1572) and [U-^2^H]betaine (DLM-407-PK) were purchased from Cambridge Isotope Laboratories and stored in stocks of 10 mg/mL at −20^°^C. All primers were synthesized by IDT (Coralville, IA). Primers used for sequencing (FWD/REV): MTR-1 = (CCCCAAAGGACACAAGGCTAA/ ACTTGCATTTTCTCCCACCAC); MTR-2 = (AGAAAGCACAGCAGTCCTGAA/ TGGATGTGCCAGCTAGTCAG). Antibodies for Western blotting were used according to their manufacturer’s directions. Anti-MTR (25896-1-AP) and Anti-COXIV (11242-1-AP) were obtained from Proteintech. All other standard laboratory chemicals used to make media were obtained from Sigma-Aldrich.

### Custom media preparation

For growth curve experiments, tracer experiments and metabolomics, cells were cultured in DMEM formulated in house from their chemical components. Standard DMEM contains folic acid (4 mg/L). For medias composed of alternative folate species (5-methyl-THF, leucovorin, or both), total folate species concentration was 4 mg/L. In addition, homocysteine (10 mg/mL) and vitamin B_12_ (1 mg/mL) were added to all custom medias. For tracer media, the amino acid found in standard DMEM was replaced with an equal concentration of tracer amino acid. For formate rescue experiments, sodium formate (141-53-7; Sigma-Aldrich) was added to a final concentration of 1 mM. For tracer, growth and metabolomic experiments, medias were supplemented with 10% (v/v) dialyzed FBS (dFBS, F0392; Sigma-Aldrich). Final media pH was adjusted to ~7.4 prior to sterile filtration using a Stericup Sterile Vacuum filter (SCGPU02RE; MilliporeSigma).

### Aqueous metabolite extraction

For aqueous metabolite extraction from cultured cells, cell lines were plated and grown to 80% confluency in 6-well plates in regular DMEM supplemented with 10% FBS or in the described custom folate medias. At start of experiment, cells were washed and media replenished with that indicated (e.g. tracer media, custom folate media, formate media, etc.). After incubation for 4 h under standard conditions, media was rapidly removed by aspiration, cells were washed by PBS 3x, and metabolism was immediately quenched by the addition of 800 uL of ice cold 40:40:20 acetonitrile:methanol:water (0.5% formic acid). After 1 min of incubation on ice, the extract was neutralized by the addition of NH_4_HCO_3_ (15% w/v). Samples were incubated at −20°C for ~30 min, then scraped, transferred to Eppendorf tubes, and centrifuged (15min, 16000 rpm, 4 °C). The resulting supernatant was frozen on dry ice and kept at −80°C until LC-MS analysis.

For extraction of aqueous metabolites from tissues, ~10-mg tissue was disrupted using a Cryomill and lysed in 1 ml ice-cold 40:40:20 acetonitrile:methanol:water (0.5% formic acid) followed by neutralization with 15% w/v ammonium bicarbonate. Solids were precipitated and supernatants were either utilized directly for LC-MS analysis (by HILIC LC-MS) or were dried and resuspended in water (for reverse phase LC-MS).

### Folate metabolite extraction

Folate species were isolated using a previously established protocol [53]. Briefly, cells in culture were seeded at 5×10^6^ cells/well in triplicate in 6-well plates maintained for at least 2 doublings in their designated custom folate medias. After aspirating media and washing 3x with PBS, metabolism was quenched with ice-cold extraction buffer (50:50 H_2_O:MeOH, 25 mM sodium ascorbate, 25 mM NH_4_OAc, pH 7). For xenografts, tumors were weighed and 50 mg of tissue was disrupted via cryomilling. The tissue was then extracted in 1 mL ice-cold buffer (50:50 H_2_O:MeOH, 25 mM sodium ascorbate, 25 mM NH_4_OAc, pH 7). For both cells and tumors, solids were precipitated by centrifuging at 16000g at 4°C for 10 min. The resulting supernatant was dried under N_2_ flow and resuspended in a 50 mM K_3_PO_4_ solution (30 mM ascorbic acid, 0.5% 2-mercaptoethanol, pH 7). Rat serum (R9759, Sigma-Aldrich) was added and samples incubated at 37°C for 2 hours to cleave polyglutamate tails on folate species. Sample pH was then adjusted to ~4 with formic acid prior to loading on Bond Elut-PH SPE columns (Agilent, 14102005), washing with aqueous buffer (30 mM ascorbic acid, 25 mM NH_4_OAc, pH 4.0), and eluting with 50:50 H_2_O:MeOH (0.5% 2-mercaptoethanol, 25 mM NH_4_OAc, pH 7). Subsequent eluate was dried down under N_2_ flow, resuspended in HPLC-grade water, and centrifuged to remove insolubles. The supernatant was then analyzed immediately or stored at −80°C until LC-MS analysis.

### Generation of MTR knockout cell lines

Generation of batch MTR knockout cells in each cell line was achieved using the lentiviral CRISPR–Cas9 vector system LentiCRISPR v2 (Addgene #52961). Briefly, RNA sequences targeting exon 15 and exon 28 of human MTR were designed using the crispr.mit.edu design tool. The guide RNA or scrambled control sequences (Supplementary Table S1) were subcloned into the lentiCRISPR v2 using the BsmBI restriction endonuclease (NEB R0580S). Virus was produced through PEI (MilliporeSigma, 408727) transfection of vectors and lentiviral packaging plasmids psPax2 and VSVG in 293T cells. Medium containing lentivirus was collected after two days and filtered through a PES filter (0.22 μm, MilliporeSigma). Cells were transfected with virus containing scrambled control or targeting *MTR* and Polybrene (8 ug/mL, Invitrogen). Cells were split after 48 h into standard RPMI-media (10% FBS) containing puromycin (2 ug/mL) and cultured for 3 days. The resulting batch knockout cells were suspended in RPMI-media (20% FBS) and plated at a dilution of ~1 cell per well in 96-well plates. After 10 days, single colonies were identified and passaged. MTR knockout of clonally isolated knockout cells was confirmed by Sanger sequencing and by western blotting.

### Cell growth experiments

Media containing different folate species were formulated in house from their chemical components. Clonally isolated MTR knockout cells in the HCT116, 8988T, and HepG2 backgrounds were plated (1 μL of cells per well) in 6-well plates in the indicated conditions. When approaching confluency (3-5 days depending on cell line), cells were passaged and re-plated at the same dilution (1 μL of cells per well). Growth was monitored at each point by packed cell volume (PCV). If total well PCV reached 0.5 μL/mL or lower, the measurements were terminated for that condition.

### Subcutaneous xenograft growth studies

Animal studies were approved and conducted under the supervision of the Princeton University Institutional Animal Care and Use Committee. For tumor growth studies, 7-wk-old female CD1/nude mice were bilaterally injected in the rear flanks with either HCT-116 wild-type, control-1, or ΔMTR-1/ 2 cells (1×10^6^ cells in 100 μL 1:1 PBS:Matrigel). Tumor growth measurements were taken biweekly using two caliper measurements [volume=1∕2 (length×width^2^)]. Animals were euthanized when tumors reached ~1500 mm^3^ or if they displayed any signs of distress or morbidity. Upon termination of the study, tumors were isolated, weighed and snap-frozen in liquid nitrogen using a Wollenberger clamp. Tumor tissues were stored at −80^°^C prior to analysis.

### Mouse infusion studies

Animal studies were approved and conducted under the supervision of the Princeton University Institutional Animal Care and Use Committee. PDAC tumor tissues derived from LSL-Kras^G12D/+^; LSL-Trp53^R172H/+^; and Pdx-1-Cre (KPC) mice were used to generate a syngeneic xenograft model, as previously described [38]. Briefly, tumor tissues were minced, mixed with Matrigel (Corning Cat #354234 in 1:1 ratio (v/v)) and injected (200 μL) into the subcutaneous left flank of C57BL/6J mice (Charles River). Tumors were allowed to grow until palpable, at which point jugular vein catheter placement surgery was performed. Mice were single-housed and allowed to recover for 7 days following surgery. On day of infusion, food was removed ~6 h prior to start. Isotope tracers in saline ([U-^13^C]methionine, 40 mM, 0.1 μl/g/min; [U-^13^C]serine, 50 mM, 0.1 μl/g/min) were infused via the catheter. After 2.5 h, serum was collected via the tail vein. Mice were euthanized via cervical dislocation and tissues (including tumor) were isolated immediately and snap-frozen in liquid nitrogen using a Wollenberger clamp to preserve metabolite levels. For betaine infusions [U-^2^H]betaine (50 mM, 0.1 μl/g/min), pre-catheterized, non-tumor bearing C57BL/6J mice from Charles River were utilized and infused for 4 h. All tissues were snap frozen using a Wollenberger clamp and stored at −80^°^C prior to analysis.

### LC-MS Analysis

Aqueous metabolites were analyzed using a quadrupole-orbitrap mass spectrometer (Q Exactive Plus, Thermo Fisher Scientific, Waltham, MA), coupled to hydrophilic interaction chromatography (HILIC) with LC separation on a XBridge BEH Amide column (Waters), and a stand-alone orbitrap (Thermo-Fisher Exactive) coupled to reversed-phase ion-pairing chromatography with LC separation on a HSS-T3 column (Waters). Both mass spectrometers were coupled to their respective liquid chromatography methods via electrospray-ionization and were operating in negative ion mode with a scan range m/z 75-1000. Detailed analytical conditions have been previously described [54, 55]. Betaine measurements were made using a Q Exactive Plus operating in positive ion mode with a scan range m/z 75-1000 coupled via electrospray-ionization to reverse-phase chromatography with LC separation on an Agilent Poroshell 120 Bonus-RP column [56]. Folate extracts were analyzed by Q Exactive Plus operating in negative ion mode with a scan range m/z 350-1000, coupled via electrospray-ionization to reverse-phase chromatography with LC separation on an Agilent Poroshell 120 Bonus-RP column. Cell culture metabolite abundances were normalized by packed cell volume; tissues and tumor metabolite abundances were normalized by mass. Isotopic labeling of metabolites arising from ^13^C-labeled nutrients were corrected for natural abundance [57]. All LC-MS data was analyzed using the ElMaven software (v 0.2.4, Elucidata), with compounds identified based on exact mass and retention time matched to commercial standards.

### Statistics

Samples sizes are defined in each figure legend. Results for biological replicates are presented as mean ± SD or mean ± SEM, as noted in figure legends. Statistical significance was calculated using unpaired Student’s t-test when comparing two groups, and one-way ANOVA followed by Dunnett’s post hoc analysis when comparing three or more. All statistical calculations were performed using the software package GraphPad Prism 7.03.

## Acknowledgments

We thank Wenyun Lu and other members of the Rabinowitz laboratory for helpful comments and suggestions. LentiCRISPR v2 was a gift from Feng Zhang (Addgene plasmid # 52961). This work was supported by NIH grants 1DP1DK113643 and R01 CA163591 to J.D.R.

## Author contributions

J.M.G. conceived the study. J.M.G., J.Z.W., and J.D.R. designed the experiments. J.Z.W., J.M.G., and L.Y. conducted the experiments. J.Z.W., J.M.G., and J.D.R. wrote the paper with input from L.Y.

