## Supplemental Table and Figures for "Methionine synthase supports tumor tetrahydrofolate pools"

### Supplementary Figure Legends

#### Supplementary figure S1.

(*A*) Schematic of methionine labeling from [U-$^{13}$C]methionine. Red circles indicate $^{13}$C tracer. MTR = methionine synthase. (*B*) Schematic of methionine labeling from [U-$^{13}$C]- or [3-$^{13}$C]serine. Blue circles indicate $^{13}$C tracer. MTHFR = methylenetetrahydrofolate reductase, SHMT = serine hydroxymethyltransferase. (*C*) Methionine labeling in cell lines after culturing for 4 h in media containing [U-$^{13}$C]serine (for 293T) or [3-$^{13}$C]serine (for HCT116 and HepG2) (mean ± SD, n = 2 for HCT116 cells, 3 for 293T and HepG2). Labeling of (*D*) serine and (*E*) methionine in serum, PDAC tumors, and normal tissues of C57BL/6J mice after [U-$^{13}$C]serine infusion for 2.5 h. (mean ± SD, n=3 mice; two technical replicates were included for each tumor). (*F*) Schematic of methionine labeling from [U-$^{2}$H]betaine. Orange circles indicate $^{2}$H tracer. BHMT = betaine-homocysteine S-methyltransferase, DMG = dimethylglycine. (*G*) Betaine labeling (M+11) in normal tissues in C57BL/6J mice after [U-$^{2}$H]betaine infusion for 4 h (mean ± SD, n=1; two technical replicates were included for each tissue).

#### Supplementary figure S2.

(*A*) Expression of MTR in the HCT116, 8988T, and HepG2 cell lines as reported in the Cancer Cell Line Encyclopedia[58]. (*B*) Cell growth curves in the media containing indicated folate sources (mean ± SD, n=2). (*C*) Individual tumor volumes for HCT116 xenografts. (*D*) Terminal tumor mass of HCT116 xenografts (mean ± SEM, n=10). * = $p < 0.05$, *** = $p < 0.001$ for comparison to control.

#### Supplementary figure S3.

(*A*) Water-soluble metabolite levels from HCT116 control and MTR knockout cells cultured in indicated media conditions. Each box reflects one independent biological measurement, normalized to the average of control cells cultured in folic acid. (*B*) Relative nucleotide mono- and diphosphate abundances in HCT116 control and MTR knockout cells in indicated medias. Intensities are normalized to the average of control-1 cells in folic acid (mean ± SD, n=3). (*C*)

Relative thymidylate species abundances in HCT116 control and knockout cells in indicated medias. Intensities are normalized to the average of control-1 cells in folic acid (mean ± SD, n=3). TIC = total ion count, * = $p < 0.05$, ** = $p < 0.01$, *** = $p < 0.001$.

**Supplementary figure S4.**

(*A*) Water-soluble metabolites levels from HCT116 control and MTR knockout subcutaneous tumors. For each condition, individual biological replicates are shown, normalized to wild-type tumors and analyzed in parallel by LC-MS. WT = wild-type.

#### Supplementary Table S1.

| Name | Gene | Exon | Targeting Sequence | Forward Primer | Reverse Primer |
|---|---|---|---|---|---|
| Control-1 | scrambled | -- | ACGGAGGCTAAGCGTCGCAA | CACCGACGGAGGCTAAGCGTCGCAA | AAACTTGCGACGCTTAGCCTCCGT |
| ΔMTR-1 | *MTR* | 15 | TTGGAGGAGTCGATGCACAA**AGG** | CACCGTTGGAGGAGTCGATGCACAA | AAACTTGTGCATCGACTCCTCCAAC |
| ΔMTR-2 | *MTR* | 28 | ACCTGGGTCCCAATAAACGT**GGG** | CACCGACCTGGGTCCCAATAAACGT | AAACACGTTTATTGGGACCCAGGTC |

Supplementary figure S1.

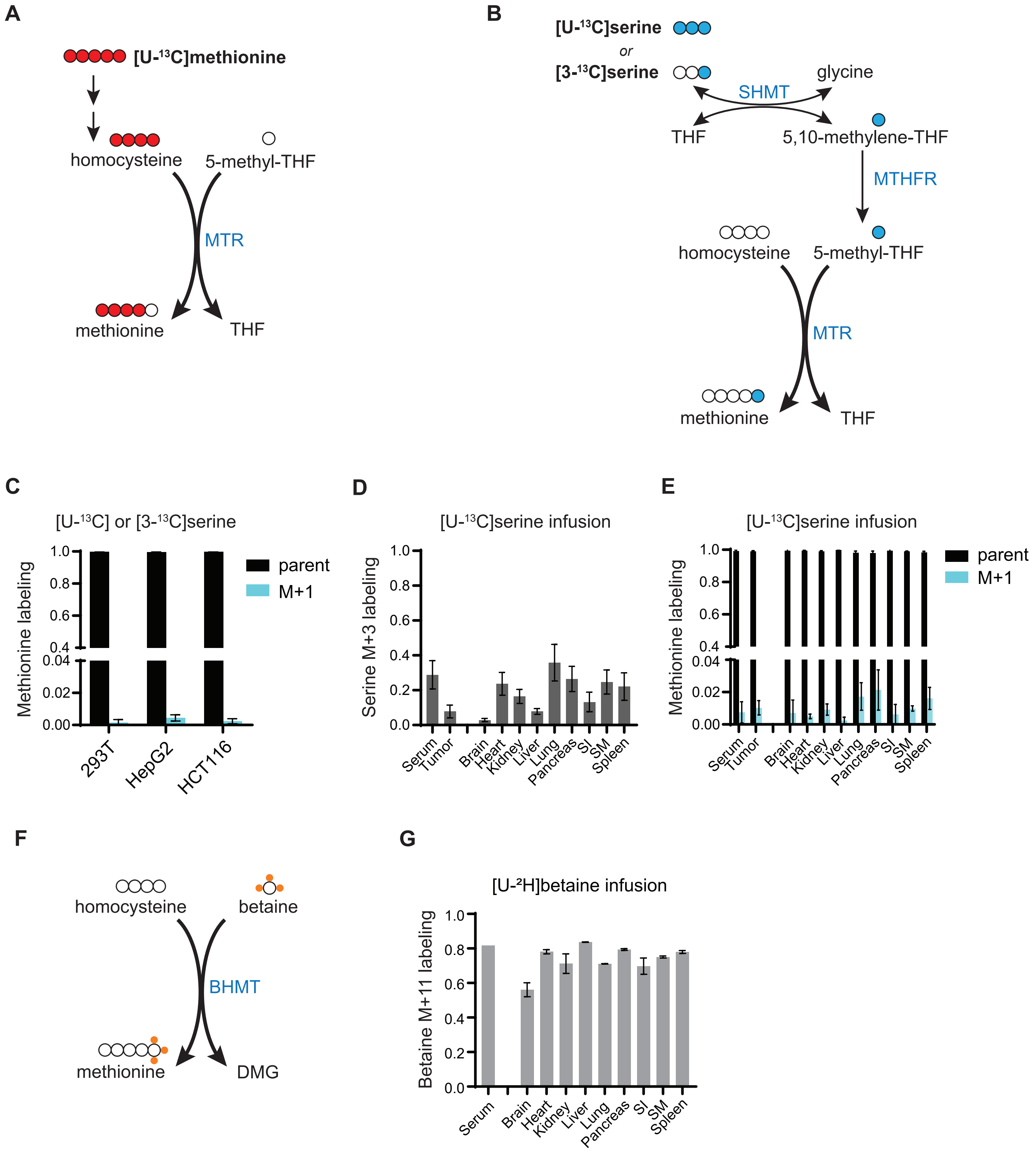

Supplementary figure S2.

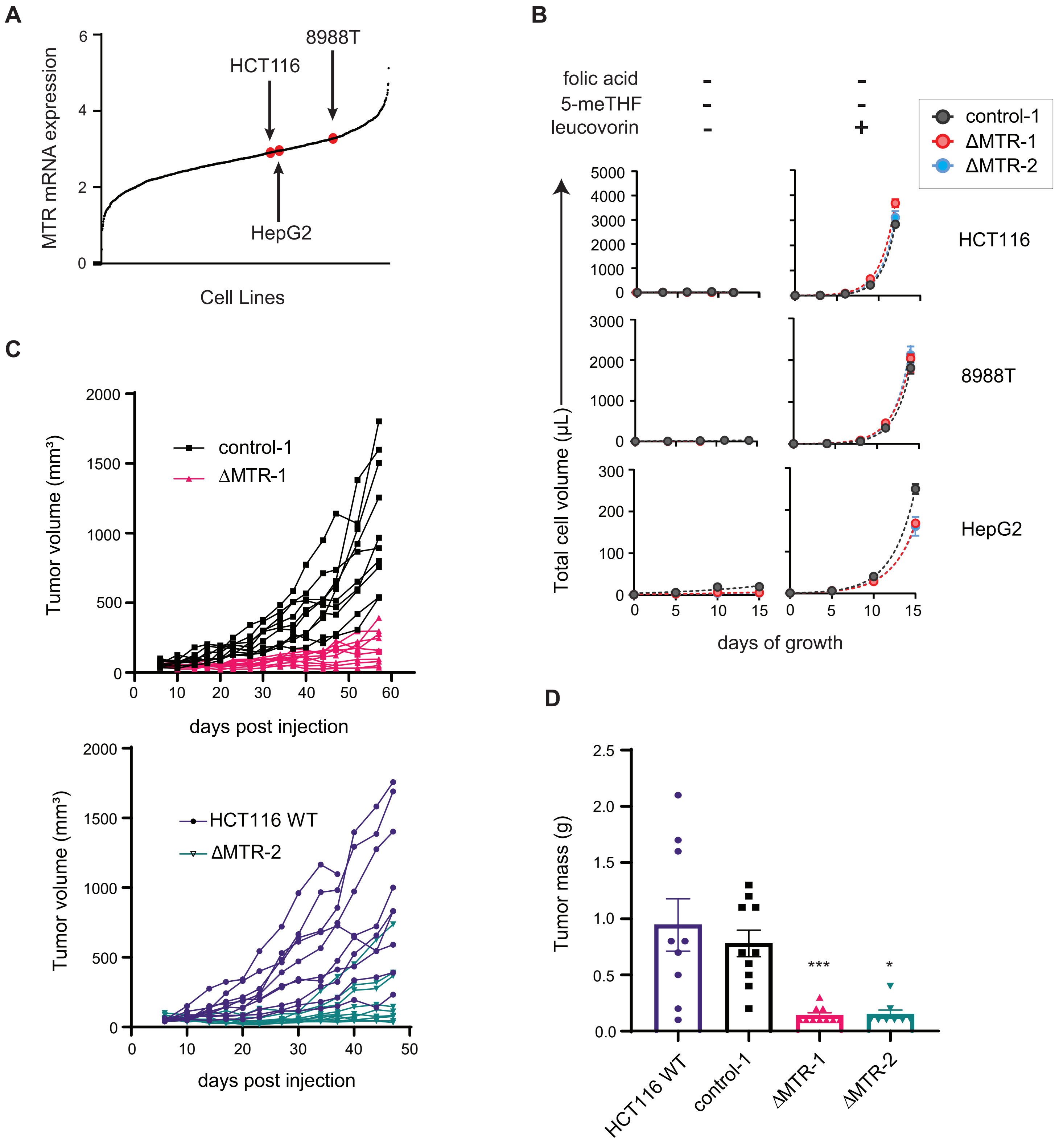

Supplementary figure S3.

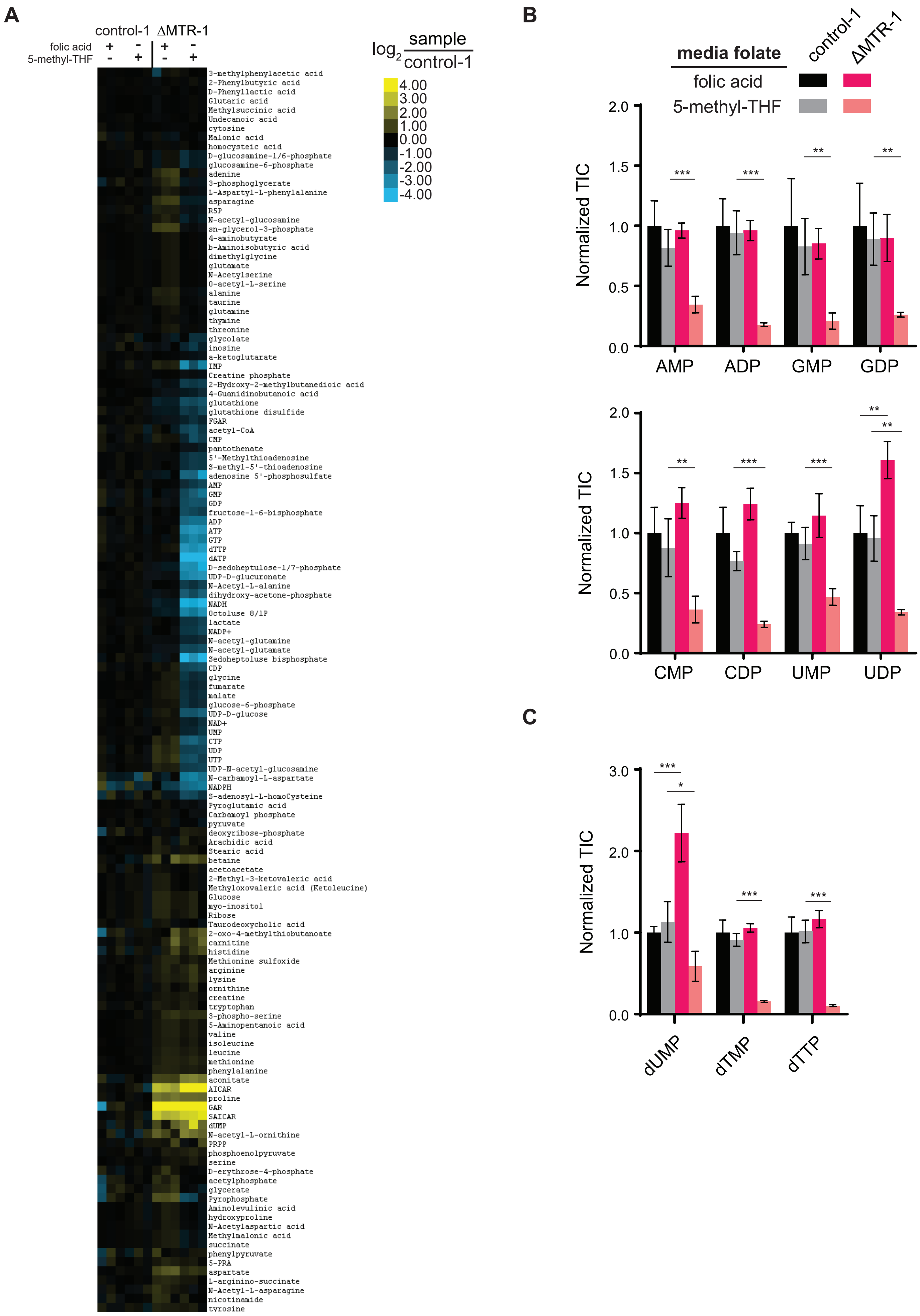

Supplementary figure S4.

**A**

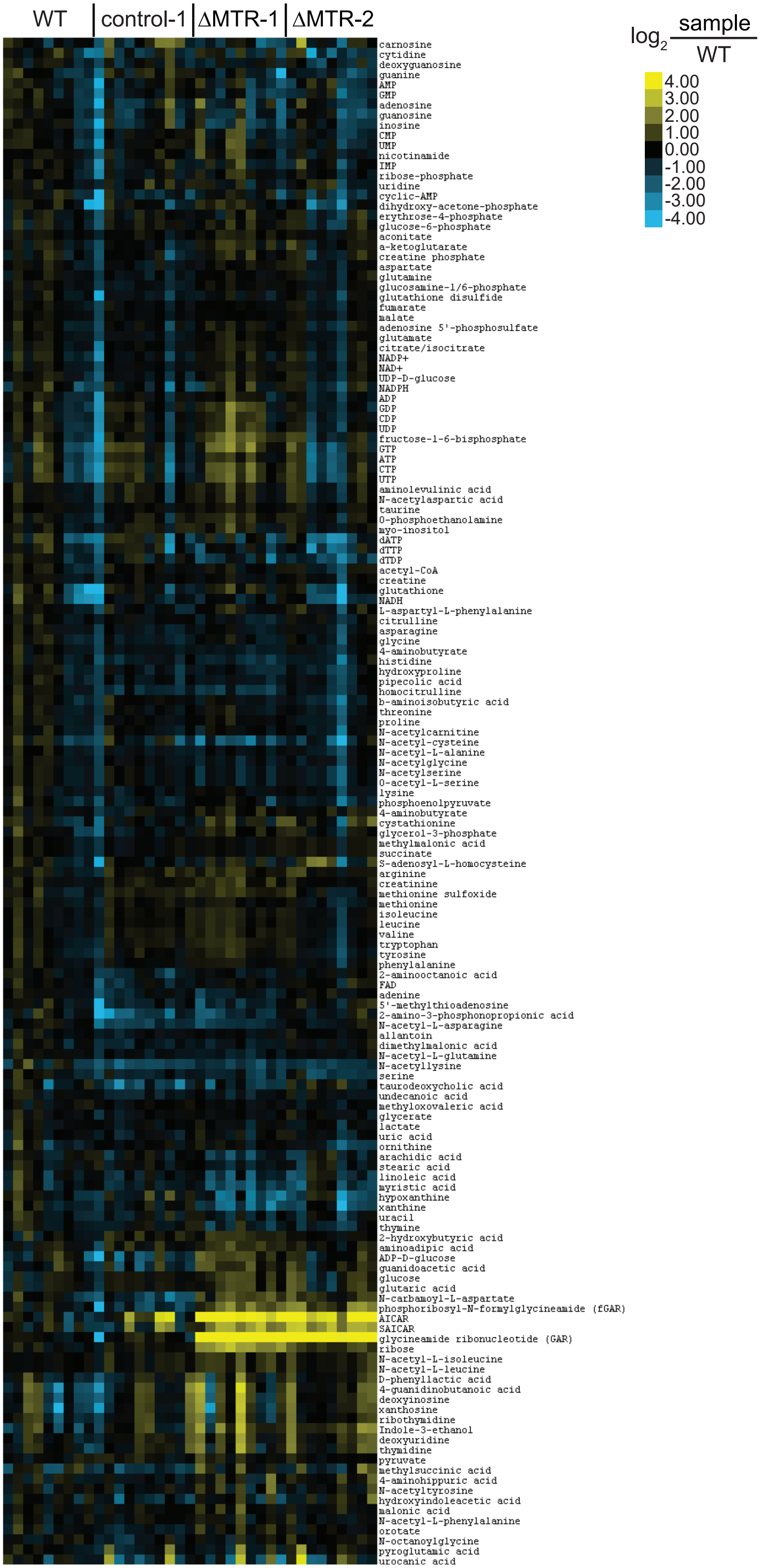
